# The Updated Mouse Universal Genotyping Array Bioinformatic Pipeline Improves Genetic QC in Laboratory Mice

**DOI:** 10.1101/2024.02.29.582794

**Authors:** Matthew W Blanchard, John Sebastian Sigmon, Jennifer Brennan, Chidima Ahulamibe, Michelle E Allen, Ralph S Baric, Timothy A Bell, Joeseph Farrington, Dominic Ciavatta, Marta Cruz Cisneros, Madison Drushal, Martin T Ferris, Rebecca Fry, Christiann Gaines, Bin Gu, Mark T Heise, Richard Austin Hodges, Tal Kafri, Rachel Lynch, Terry Magnuson, Darla Miller, Caroline E Y Murphy, David Truong Nguyen, Kelsey E Noll, Megan Proulx, Chris Sassetti, Ginger D Shaw, Jeremy M Simon, Clare Smith, Myrek Styblo, Lisa Tarantino, Joyce Woo, Fernando Pardo Manuel de Villena

**Affiliations:** Department of Genetics, University of North Carolina, Chapel Hill, North Carolina, 27599, USA; Mutant Mouse Resource and Research Center, University of North Carolina, Chapel Hill, North Carolina, 27599, USA; Department of Computer Science, University of North Carolina, Chapel Hill, North Carolina, 27599, USA; Systems Genetics Core Facility, University of North Carolina, Chapel Hill, North Carolina, 27599, USA; Department of Epidemiology, Gillings School of Public Health, University of North Carolina, Chapel Hill, North Carolina, 27599, USA; Department of Environmental Sciences and Engineering, Gillings School of Public Health, University of North Carolina, Chapel Hill, North Carolina, 27599, USA; Department of Neuroscience, The Ohio State University; Department of Microbiology and Immunology, University of North Carolina, Chapel Hill, North Carolina, 27599, USA; Lineberger Comprehensive Cancer Center, University of North Carolina, Chapel Hill, North Carolina, 27599, USA; Department of Microbiology and Physiological Systems, UMass Chan Medical School; Department of Molecular Genetics and Microbiology, Duke University; Department of Nutrition, Gillings School of Public Health, University of North Carolina, Chapel Hill, North Carolina, 27599, USA

**Keywords:** genetic QC, genetic background, substrains, chromosomal sex, genetic constructs, diagnostic SNPs, microarrays, inbred strains

## Abstract

The MiniMUGA genotyping array is a popular tool for genetic QC of laboratory mice and genotyping of samples from most types of experimental crosses involving laboratory strains, particularly for reduced complexity crosses. The content of the production version of the MiniMUGA array is fixed; however, there is the opportunity to improve array’s performance and the associated report’s usefulness by leveraging thousands of samples genotyped since the initial description of MiniMUGA in 2020. Here we report our efforts to update and improve marker annotation, increase the number and the reliability of the consensus genotypes for inbred strains and increase the number of constructs that can reliably be detected with MiniMUGA. In addition, we have implemented key changes in the informatics pipeline to identify and quantify the contribution of specific genetic backgrounds to the makeup of a given sample, remove arbitrary thresholds, include the Y Chromosome and mitochondrial genome in the ideogram, and improve robust detection of the presence of commercially available substrains based on diagnostic alleles. Finally, we have made changes to the layout of the report, to simplify the interpretation and completeness of the analysis and added a table summarizing the ideogram. We believe that these changes will be of general interest to the mouse research community and will be instrumental in our goal of improving the rigor and reproducibility of mouse-based biomedical research.

## INTRODUCTION

Genotyping arrays have been a staple in mouse research for more than two decades and have been successfully adopted for genetic QC and colony maintenance (Petkov et al. 2004; Yang et al. 2011; Morgan et al. 2015; Andrews et al. 2021; Amos-Landgraf et al. 2022). Five years after its introduction, the MiniMUGA array has been used for genotyping over 40,000 mouse samples and the manuscript describing the array and its capabilities has been cited widely (Sigmon et al. 2020; Birling et al. 2022; Bourdon and Montagutelli 2022; Yoshiki et al. 2022; Smith et al. 2022). Part of MiniMUGA’s success is due to its unique characteristics, including discrimination between commercial substrains, robust chromosomal sex determination, and detection of commonly used constructs. In addition, a key advantage of MiniMUGA is the inclusion of a descriptive summary report of the sample that accompanies cost-effective genotypes. This has made MiniMUGA an attractive tool for organizations charged with genetic quality control (GQC) of important collections such as the Mutant Mouse Resource and Research Centers (Amos-Landgraf et al. 2022, https://www.mmrrc.org/), and of large and complex breeding programs like the Collaborative Cross (https://csbio.unc.edu/CCstatus/index.py). Despite these successes and the fact that the MiniMUGA content is now fixed (i.e. production array), several limitations of the original analysis pipeline can be addressed now. These limitations were caused by the limited number and type of samples genotyped in the production version of the array at the time (less than 1,500 out of almost 8,000 used in that publication) and used for the initial marker annotation and bioinformatic pipeline validation (Sigmon et al. 2020). This had severe consequences in marker annotation from performance to diagnostic information. Additional limitations included the exclusive use of a “greedy” algorithm for determining the contribution of primary and secondary genetic backgrounds, the exclusion of Y chromosome and mitochondria from the ideogram, and the use of an overly restrictive threshold for the level of sample inbreeding required to run the informatic pipeline to completion.

Here, we report our efforts to improve marker annotation based on over 8,500 samples genotyped in the final array, improvement in the analysis pipeline for background selection, ideogram content, construct detection, and a new section that summarizes the genome complement in the form of diplotype intervals. These changes improve the rigor of the analysis and make the report useful for non-inbred samples such as experimental crosses and partially congenic lines. We are committed to continuing this cycle of improvements in the future as needed.

## MATERIALS AND METHODS

### Mice

We used 8,559 mouse DNA samples to analyze and annotate SNP marker performance. Samples included in this set were required to have excellent genotyping quality on the production version of the MiniMUGA array. In addition, we selected samples with varying levels of inbreeding and different genetic backgrounds to increase the likelihood of capturing all three possible genotypes at each SNP marker (reference, alternative, and heterozygous (hereafter ref, alt, and het respectively)). We used 1,623 mouse DNA samples from 237 distinct inbred strains to create new consensus reference genotypes. 1,011 of these consensus samples were included in the previously described marker performance sample set. 612 consensus samples were not included in the performance sample set, 352 genotyped on the initial version of the array, 236 biological replicates of CC strains already represented in the performance set, and 24 technical replicates. Eighty-three were provided by Transnetyx and Neogen to represent six additional strains and to increase the number of biological replicates in three existing strains. We used six mice provided by the MMRRC as positive controls for the Flp construct and four mice included in the initial description of the array as positive controls for the cHS4 construct. **Table S1** lists the origin of the samples included in the SNP marker performance annotation, expanded consensus genotypes, and expanded number of constructs detected. Samples are identified with random six-digit ID and a reference of the publication if available.

An additional 16,123 samples genotyped by the FPMV lab at UNC have been used for manual curation and validation of our results.

### Marker Annotation

The updated marker annotation is provided in **Table S2**. This file includes marker annotations for the following fields (fields in **bold** are new or have updated information):

- chromosome – Marker chromosomal location. Possible values are 1-19 autosomes, X and Y sex chromosomes, PAR pseudoautosomal region, or 0 for unmapped genetic construct probes.
- position – location in base pairs based on the GRCm38 reference build.
- name – marker name
- **tier_2022** – new SNP marker performance tier assignment. 1A, 2A, 1B, 2B, 1C, 2C, and 4 are possible values. Construct probes have no tier_2022 assignment.
- **diagnostic** – the list of substrains (or diagnostic class) for which this SNP is found to be diagnostic, or blank if it is not diagnostic.
- **partial_diagnostic** - the list of substrains for which this SNP is found to be partially diagnostic, or blank if it is not partially diagnostic.
- **diagnostic_allele** – the minor (diagnostic) allele at this diagnostic SNP.
- **construct_info** (called diagnostic_info in the code) – this field indicates the genetic construct this marker detects. This field is blank if the marker does not detect a construct.
- **positive_threshold** – for a set of markers that detect a given genetic construct, this value indicates the minimum threshold for reporting a positive detection (presence), based on the sum intensity of that set of markers.
- **negative_threshold** – for a set of markers that detect a given genetic construct, this value indicates the maximum threshold for reporting a negative detection (absence), based on the sum intensity of that set of markers.
- **duplication** – this field indicates whether the SNP targeted by a given marker probe appears to be duplicated in the genome of some samples. Duplication may lead to a downgrade of a marker in the tier_2022 classification. The identification of duplicated SNPs is not exhaustive.
- **recluster** – Indicates whether the Illumina genotype calling parameters (clusters) were modified in this update, relative to the initial publication (Sigmon et al. 2020). After this update, Markers annotated as TRUE may generate different genotype calls for the same intensity values (or for the same sample).

### Consensus Genotypes

The updated consensus genotypes are provided in **Table S3.** This table lists inbred strains for which we have created consensus genotypes used in primary and secondary background determination and contribution.

### Synthetic Backgrounds

For each of 131 pairs of related substrains, we generate a synthetic background *in silico.* These are added to the set of consensus genotypes used in primary and secondary background determination and contribution. To generate a synthetic background, we compare the consensus genotypes at each SNP marker for a pair of substrains. If the substrain genotypes are the same call (A,T,G,C, or H), the resulting synthetic background call is that same call. If the calls are different, the resulting synthetic call is an H. Note that this is the same as assigning H calls to the synthetic background at every marker that is annotated as diagnostic for either of the substrains in the pair.

### Strain Annotation

Classical inbred strains have been assigned to one of 11 groups based on their name and origin (**Table S4).** The rows list all classical strains, while the columns list the strain groups used for the determination of the diagnosticity of SNPs and an outgroup with all classical inbred strains without diagnostic SNPs. Numbers in each cell are the number of diagnostic SNPs followed by the number of partially diagnostic SNPs, separated by a forward slash. N/A is not applicable because, by definition, they cannot have diagnostic SNPs. Blank cells should be ignored.

## RESULTS

This update of the MiniMUGA platform takes advantage of the large increase in the number of DNA samples genotyped with the production array since the original publication (Sigmon et al. 2020) to: 1) update the annotation of markers, including performance and diagnostic value for SNP markers and identification of markers used for detection of two additional constructs; 2) generate higher quality consensus genotypes for a larger number of inbred strains; and 3) update the content and layout of the report.

### Updated Annotation of SNP Marker Performance

To address the limitations of the MiniMUGA pipeline, we reassessed the performance of the 10,819 SNP markers in the array and used that information to improve their annotation. For this task, we used a set of 8,559 excellent quality samples genotyped with the production version of the array (materials and methods; annotated as PERFORMANCE sample set in **Table S1**). This set includes 1,943 inbred samples (or samples with very low levels of residual heterozygosity) and 6,616 outbred samples, including 535 F1 hybrids. Most outbred samples in this set are experimental backcrosses or intercrosses. Both sexes are similarly represented in this set, and there are a few examples of sex chromosome aneuploids (4,239 XX, 4,276 XY, 36 XO, and 8 XXY as defined by our original method (Sigmon et al. 2020). It also includes at least one sample from each inbred strain for which we have consensus genotypes. The representation of homozygous and heterozygous calls as well as both sexes is critical to improving the annotation of SNP marker performance.

We generated individual scatter plots for each SNP marker, plotting the normalized intensities for the performance sample set, using different colors to visualize the three possible genotype calls: ref, alt, and het, plus no calls (**Figure 1, Figure S1**). As expected, for most markers the allele intensities group into three distinct clusters representing the three standard types of calls (ref, alt, and het). We identified 791 markers where at least one allele cluster included multiple different genotype calls, due to incorrect genotype-calling software tuning (**Figure 1**). We used the allele intensity plots to set new cluster boundaries and recalled all 8,559 samples at these 791 markers. We then compared plots and genotype calls to identify SNPs where the new cluster boundaries produce more consistent genotype calls (fewer clusters with inconsistent genotype calls and fewer no calls overall). The new cluster boundaries were updated for 756 markers (733 from training array content, 23 from production) where there is marked improvement in the congruency of the genotype calls (compare panels **A** and **B** in **Figure 1**). Users should expect discrepancies between the new and original genotype calls at these markers, given that the cluster boundaries have changed.

**Figure 1.**
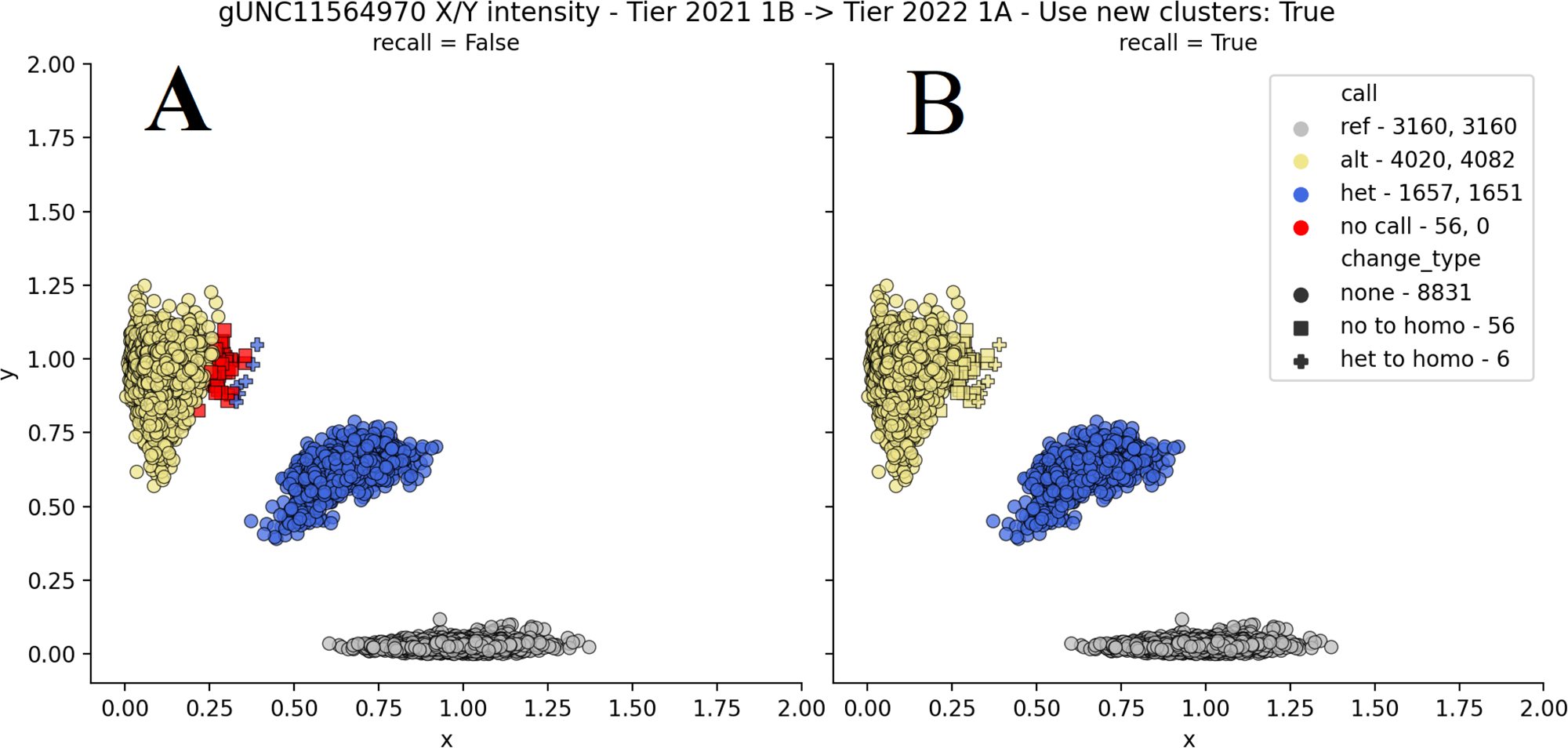
Intensity plot for Tier 1 A marker gUNC11564970. The left side shows the genotype calls for 8,893 samples with the original algorithm. Note that 6 samples were called heterozygous incorrectly (they are homozygous alt) and 56 resulted in no genotype call. These miscalls have been corrected on the right side using new clustering parameters. Each sample is shown as a circle, square, or cross. Colors denote genotype calls, while shape denotes change in genotype call.

The next step was to address the performance of individual SNPs after reclustering based on the number of clusters, the discrimination between clusters, and the level of consistency of genotype calls within a cluster in the performance sample set. We use these metrics to define a new performance annotation called Tier 2022 (henceforth called Tier). First, we divided markers into three broad classes: class 1 makers have three distinct clusters at the expected locations for reference, alternate, and heterozygous genotype calls for autosomal and X chromosome markers, and two distinct reference and alternate clusters at Y chromosome and mitochondrial markers. Class 2 markers have an additional cluster of no calls at or near the intensity plot origin (these samples fail to produce a signal for either allele). This cluster is predicted to be due to the presence of off-target variants in one or more haplotype(s) (Didion et al. 2012, **Figure S1**). We used Whole-Genome Sequencing to confirm this prediction in a small set of cases (data not shown). Tier 4 markers exhibit no recognizable intensity clustering patterns (or form only one cluster) and fail to produce reliable genotype calls in the performance sample set. These markers should always be excluded from analysis. The second level of classification applies to tier 1 and 2 markers and divides them into three additional classes, A, B, and C. Class A markers have the highest performance within a tier: the three genotype clusters are clearly separated, there are no inconsistent genotype calls within these clusters, and they have no or very few N calls, excluding the cluster of N calls at the origin for tier 2 markers. Classes B and C have decreasing performance, with lower discrimination between clusters, an increasing number of inconsistent genotype calls within clusters, and an increasing number of N calls. There is no hard boundary between B and C classes as the performance depends on the haplotype present at the locus.

Tier 1A markers have the best performance across all samples and comprise 82.5% (8,929) of all SNP markers in the array. **Table 1** summarizes the number of markers classified in each Tier. In the sample report, only Tier 1A markers are used to identify and quantify the contribution of the primary and secondary backgrounds, and to determine the level of inbreeding. Although tiers 1B, 2A and 2B may perform well in most genetic backgrounds, they are not completely reliable and may produce incorrect genotype calls in some backgrounds. For example, for tiers 2A and 2B markers, samples that are heterozygous for a combination of the off-target variant and any of the two standard alleles will genotype as homozygous. Furthermore, Tier 1B includes markers that are duplicated in the genome of one or more classical laboratory strains and may lead to heterozygous calls in some inbred mice and incorrect genotype calls in some experimental mice. Markers with known or suspected duplications in inbred strains are annotated in **Table S2**.

**Table 1.** MiniMUGA SNP Marker Counts - grouped by Tier 2022 and diagnostic capability. Each cell in the table displays the number of markers with a given Tier and Diagnostic Capability, followed by the percentage of the row total this represents in parenthesis.

| Tier 2022 |  |  |  |  |  |  |  |  |
| --- | --- | --- | --- | --- | --- | --- | --- | --- |
|  | A |  | B |  | C |  | 4 | Total |
|  | 1A | 2A | 1B | 2B | 1C | 2C |  |  |
| Diagnostic | 3098 95.2% | 106 3.3% | 44 1.4% | 5 0.2% | 0 0.0% | 0 0.0% | 0 0.0% | 3253 30.1% |
| Not Diagnostic | 5831 77.2% | 185 2.4% | 151 2.0% | 44 0.6% | 209 2.8% | 28 0.4% | 1118 14.8% | 7566 69.9% |
| Total | 8929 82.5% | 291 2.7% | 195 1.8% | 49 0.5% | 209 1.9% | 28 0.3% | 1118 10.3% | 10819 100.0% |

In addition, we manually downgraded the performance annotation of specific makers if we observed incorrect genotype calls in samples with known genotypes (inbred, F1 and F2 mice).

### Improved Annotation of Diagnostic SNPs and Alleles

We classified each classical inbred strain into one of ten strain groups or a common outgroup (see **Table S4**). The classification was based on partial name-sharing and historical records. A given marker is considered diagnostic if its minor allele is present in only one strain group and absent in the common outgroup. For these diagnostic markers, the set of substrains in which the minor (now diagnostic) allele is observed is called its diagnostic class.

Diagnostic alleles observed in some but not all constituent samples used to create the genotype consensus for a given substrain are annotated as “partially diagnostic”. Note that partially diagnostic SNPs are typically not shared between substrains. Partially diagnostic SNPs are annotated as heterozygous (or H) in the consensus for the substrain in which they are segregating (**Table S3**).

By design, there is an overrepresentation (3,253) of diagnostic SNPs in MiniMUGA to ensure that it can discriminate between as many substrains as possible. The performance of diagnostic SNPs is, on average, better than non-diagnostic markers. For example, 95% of diagnostic markers are in Tier 1A versus 77% of non-diagnostic markers. In addition, there are no diagnostic markers in Tier 4 versus 15% of non-diagnostic markers (**Table 1**). In conclusion, the presence of diagnostic alleles in a sample is highly reliable evidence for the contribution of the corresponding substrain(s) to the genome of the sample.

### Expanded Detection of Genetic Constructs

We annotated 14 probes (**Figure S2**) which detect two additional constructs: chicken HS4 insulator (cHS4) and Flippase (Flp). **Figure 2** shows the aggregate performance of these probes in negative controls, positive controls, and experimental samples. Thresholds for presence and absence were determined as previously described for other constructs by minimizing the number of experimental samples with questionable presence of the corresponding constructs while keeping positive and negative controls fully concordant (Sigmon et al. 2020).

**Figure 2.**
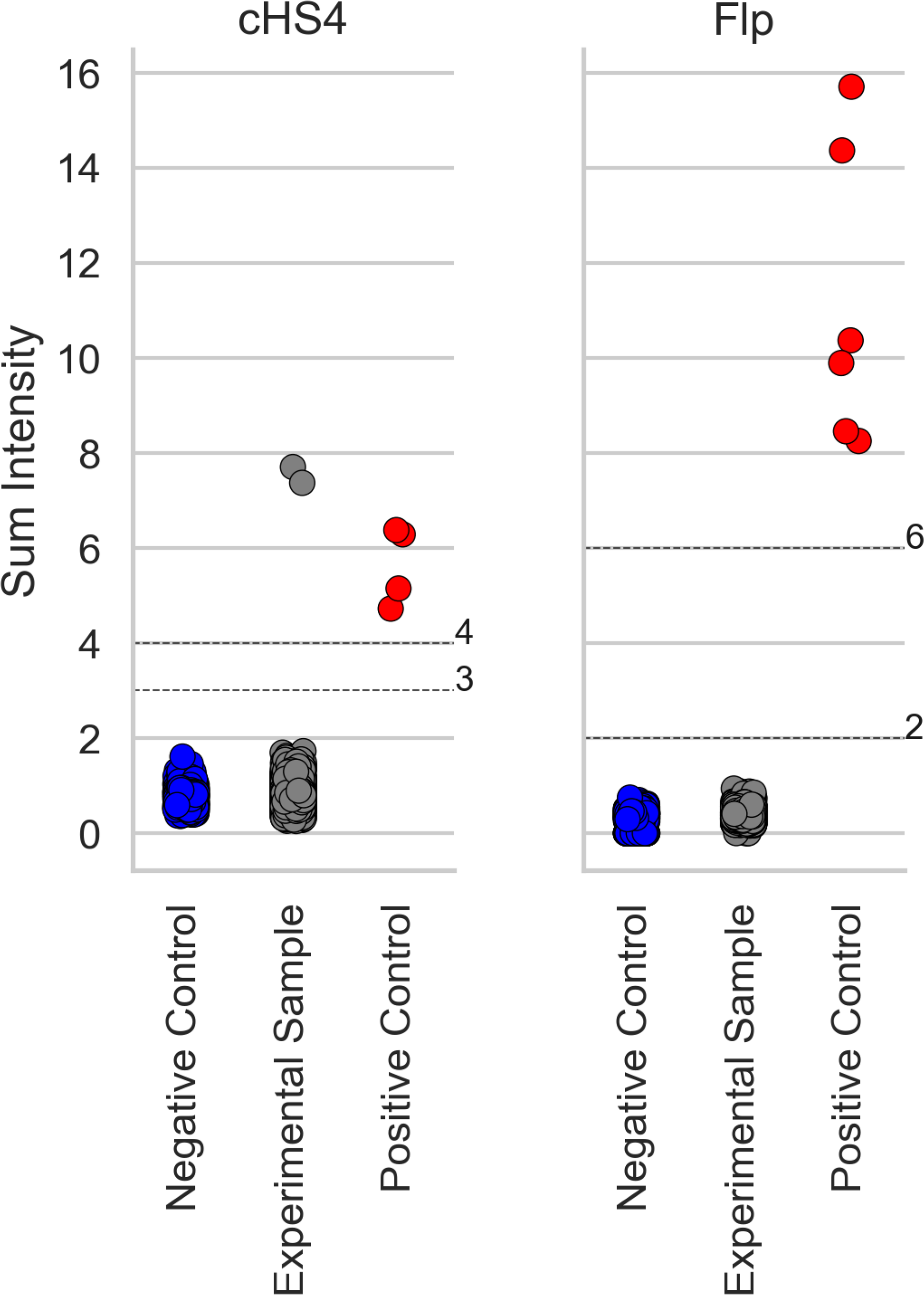
Detection of new genetic constructs validated in MiniMUGA. For each construct, samples are shown as dots and classified as negative controls (left), experimental (center), and positive controls (right). The dot color denotes whether the sample is determined to be negative (blue), positive (red), or questionable (gray) for the respective construct. For each construct, the gray horizontal lines represent data-driven ad hoc thresholds discriminating between presence and absence. Chicken HS4 insulator (cHS4) and Flippase (Flp).

### Expansion and Update of Consensus Genotypes for Inbred Strains

We created new consensus genotypes for 237 inbred strains based on 1,623 samples (only 556 samples were used to create the previous consensus genotypes). All samples in the consensus set were curated to ensure they were pure representatives of the corresponding inbred strain. This consensus set includes five additional classical inbred strains (BALB/cAnNRj, BALB/cByJRj, BALB/cJRj, CBA/CaJ, DBA/1Rj) and updated genotypes for 63 Collaborative Cross (CC) strains.

Consensus genotype calls are a representation of the strain or substrain genotypes. The accuracy of a given strain consensus depends on the number of biological replicates included. The calls are generated according to previously described rules (Sigmon et al. 2020), with one notable change: partially diagnostic SNPs, which were previously annotated as a lower-case diagnostic allele call, are now annotated as H in the corresponding substrain. This H call indicates that the diagnostic allele is segregating in a substrain; therefore, samples from this substrain may have any of the three possible genotypes at those markers.

**Figure 3** shows the distribution of biological replicates per strain, grouped by strain type (**Table 2**). The number of replicates is variable. The average number of replicates in the 89 classical inbred strains is 5.76 (range 1-28). For classical inbred strains where we have multiple substrains or sibling strains, there is a higher number of replicates. The number of replicates for wild-derived stains is lower than for classical inbred strains. For the CC strains, the average number of samples in the consensus is higher (15.06), it always includes males and females, and includes all breeders alive for that strain in 2020 in the SGCF at UNC. The BXD panel typically includes only one sample per strain.

**Figure 3.**
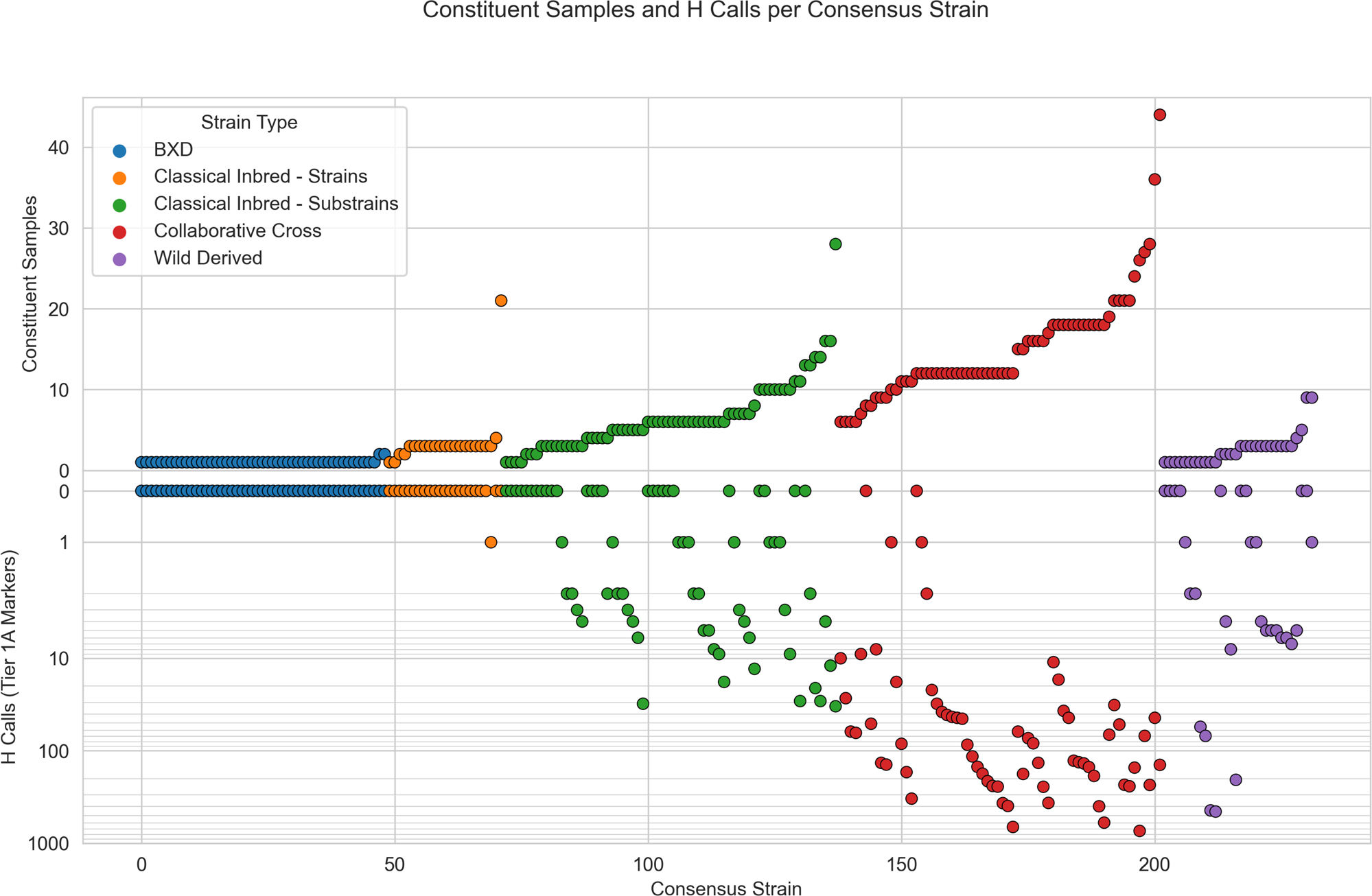
Number of biological replicates and heterozygous calls in the consensus genotypes of each inbred strain. Each dot is an inbred strain, color coded into five groups according to the legend. Classical inbred strains are divided into two groups based on the fact that they are part of a group of substrains or sibling strains (diagnostic, green) or independent inbred strains (orange). Strains are ordered by ascending number of replicates within type. Only Tier 1A markers are included in the analysis. The upper Y axis is the number of replicates per consensus strain. The lower Y axis is the number of H calls per consensus strain, in an inverted log10 scale.

**Table 2.** Constituent samples per consensus strain.

| Strain Type | strains | Total Samples Per Strain |  |  | Male Samples Per Strain |  |  |
| --- | --- | --- | --- | --- | --- | --- | --- |
|  |  | mean | min | max | mean | min | max |
| Classical Laboratory | 89 | 5.83 | 1 | 28 | 3.81 | 0 | 15 |
| Wild Derived | 34 | 2.53 | 1 | 9 | 2.18 | 0 | 5 |
| CC | 64 | 15.06 | 6 | 44 | 5.14 | 2 | 13 |
| BXD | 49 | 1.04 | 1 | 2 | 0 | 0 | 0 |
| Other Recombinant Inbred | 1 | 3 | 3 | 3 | 3 | 3 | 3 |

**Figure 3** also shows the number of consensus H calls per strain, below the X-axis. A priori, inbred strains should have no H calls for tier 1A markers. However, H calls can represent either true heterozygosity (in partially diagnostic markers for substrains and residual heterozygosity in the CC strains) or rare miscalls due to off-target variants. The latter are particularly prevalent in wild-derived strains (note that the CC strains are derived from eight founders including three wild-derived strains).

### Changes in the Analytical Pipeline and Report Layout

There are five main updates in the analysis pipeline and/or report presentation (**Figure 4**): 1) updated pipeline for genetic background determination in samples with more than one genetic background; 2) changes in inbreeding estimation; 3) inclusion of the Y Chromosome and mitochondria in the ideogram; 4) specification of the ‘Minimal Strain Sets Explaining All Diagnostic Classes’ and 5) the addition of a table with diplotype intervals.

**Figure 4.**
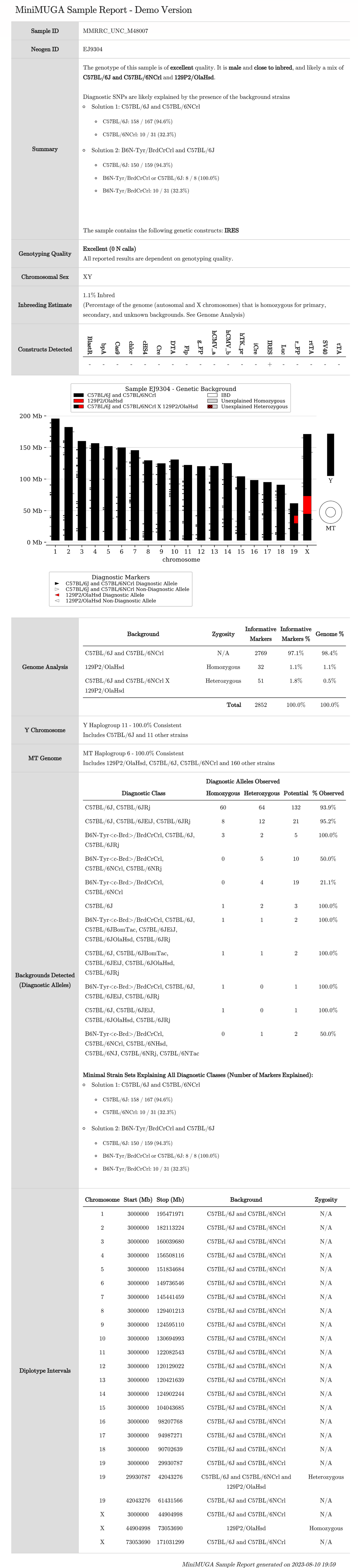
The figure is an example of the current layout of the MiniMUGA report for an MMRRC congenic sample.

### Determination of Global Genetic Background

The Genome Analysis section summarizes the global genetic background, showing the contribution of the primary, secondary, and further unexplained backgrounds, each as a percentage of the genome. This global summary complements the local information shown in the ideogram. The primary and secondary backgrounds are determined by the greedy algorithm described previously (Sigmon et al. 2020) with the following changes: 1) Background analysis now involves only the highest quality markers (Tier 1A); 2) The set of potential background strains now includes several synthetic backgrounds composed of pairs of substrains generated *in silico* (see Materials and Methods). These were created specifically to enable the pipeline to work equally well for samples whose genetic background includes two related substrains plus an additional third (or even a fourth) strain; 3) The default threshold for clustering remains set at two markers per 2Mb; 4) The requirement for samples to have less than a threshold level of heterozygosity has been removed and thus reports are created for all samples including F1 hybrids and F2 samples. As a result, the updated pipeline produces a complete report at more than twice the rate of the original (only 11,767 samples out of 26,245 (44.8%) in our database would have generated a full report with the original pipeline); and 5) The threshold for the minimum fraction of the genome that must be explained to produce a report is now 97.5% percent (99.8% previously).

In cases where two backgrounds are necessary to explain a sample, we only use markers that are informative among the primary background, secondary background, and sample genotypes. We use the genome positions of these informative SNPs to identify clusters in the sample genome of consistent genetic background and zygosity. We merge neighboring clusters with the same background and zygosity. We extend the proximal and distal cluster boundaries to the midpoint for the intervals between clusters with different backgrounds and/or zygosity. For the most proximal and distal clusters, we extend them to the start and end of the corresponding chromosome. The resulting set of Diplotype Intervals is used to construct the Ideogram presented in the Local Genetic Background section of the report, to estimate the contribution of the primary, secondary, and unexplained backgrounds presented in the Global Genetic Background, and to calculate the overall Level of Inbreeding reported for the sample. This physical size-based approach improves the estimation of the contribution of the primary and secondary backgrounds and the level of heterozygosity (or inbreeding) over the original marker number-based analysis used in the original version of the MiniMUGA sample report (Sigmon et al. 2020). In addition, the updated method of estimating the contribution of genetic background using physical distance (in megabases) is a major improvement over the original pipeline, which was based solely on the number of markers. Specifically, the initial pipeline overestimated the contribution of the primary background to the sample genome by an average of 18% (range 0-48%) (**Table S5**). It also underestimated the contribution of the secondary background in homozygosity to the sample genome by an average of 8.6% and 8.7% in homozygosity and heterozygosity respectively (ranges 0-27%, 0-33%, respectively). The contribution of any unexplained background to the genome was underestimated by 1.1% on average (0-7.1%). Error in the estimated contribution to genetic background in the original pipeline was proportional to the amount of non-primary background present in a sample. In summary, the updated pipeline performs much better than the original, particularly in samples with heterogenous genetic background, including F2 and F1 hybrids.

### Changes in the Inbreeding Estimate

The inbreeding estimate is now based on the percentage of the nuclear genome, excluding the Y chromosome, that is homozygous (or hemizygous for males) for primary, secondary, and unexplained backgrounds. In contrast to the previous estimate, this update is far less dependent on the number and density of informative SNPs for a given pair of backgrounds. This is especially relevant for samples derived from two closely related substrains (**Figure S3**).

### Representation of the Local Genetic Background in the Ideogram

The report shows the local genetic makeup of the sample in a visualization of the genome known as an ideogram. This is as described in the original paper (Sigmon et al. 2020), with the addition of the Y chromosome (shown as a single bar) and the mitochondria (shown as a torus). The regions of the genome shown in black represent the primary or majority genetic background. The regions shown in red represent the secondary or minority genetic background. Red and black in the same region indicates heterozygosity. Transitions between primary and secondary backgrounds are always placed at the midpoint between flanking informative markers (see above). Regions shown in white are Identical By Descent (or IBD). This means that MiniMUGA cannot distinguish between the primary and secondary backgrounds in these regions, and as above, this is especially relevant for samples derived from two closely related substrains. The MiniMUGA sample report is optimized for reproducible mouse models involving only one or two genetic backgrounds. If more than two backgrounds are present, any unexplained regions will be shown in gray.

The diagnostic markers for the primary and secondary background are also shown in the ideogram. Diagnostic markers for the primary background are represented by triangles at the corresponding position on the left side of the chromosome. Similarly, diagnostic markers for the secondary background are represented by triangles on the right side of the chromosome. Black-filled triangles indicate the presence of the diagnostic allele for the primary background at the corresponding SNP in the sample. Red-filled triangles indicate the presence of the secondary background at the corresponding SNP in the sample. Note that filled triangles do not imply homozygosity or heterozygosity for the diagnostic allele. Unfilled triangles indicate the absence of the diagnostic allele and can sometimes be found in unexpected regions due to the existence of partially diagnostic SNPs.

### Reporting of the Substrains Detected with the Diagnostic Alleles

Utilizing the updated diagnostic annotations, we now report the number and zygosity of SNPs with diagnostic alleles for each of the 94 diagnostic classes (**Table S2**) present in a sample. As previously discussed, a diagnostic class groups all substrains sharing specific diagnostic alleles and thus reflects the local ancestry of the substrains. Then the report programmatically determines the minimum set, or sets, of substrains required to explain all the diagnostic alleles present in a sample. This is presented as “*Minimal Strain Sets Explaining All Diagnostic Classes*” in the report. There are three important caveats to consider here. First, if there are multiple solutions presented (multiple combinations of substrains) the user is encouraged to use external information, if available, to select the most likely solution. Second, the minimal solution may not represent the genetic backgrounds present in a sample. Any combinations of substrains covering all diagnostic classes can be the “true” solution. Third, the “Minimal Strain Sets Explaining All Diagnostic Classes” is not used **directly** in the determination of primary and/or secondary background present in a sample. In other words, genotypes at those SNPs are used but not given any special weight or consideration. Finally, we reiterate the point made in the original paper (Sigmon et al. 2020) that these markers are not reliable in mice with some contribution from wild derived strains.

### Addition of Diplotype Intervals

This section presents the data shown in the ideogram in table format. The main differences with the ideogram are that diagnostic SNPs, the Y chromosome, and mitochondrial genome are not included. For each interval, the table provides the chromosome, start and end positions in GRCm38 genome coordinates, background (name(s) of the (sub)strain assigned to the interval), and zygosity. Note that for XY males, the X chromosome is reported as homozygous instead of hemizygous.

## DISCUSSION

It has been our experience since the public release of the MiniMUGA array that many users struggle with and spend an inordinate amount of effort dealing with uncommon inconsistent and/or unexpected genotype calls in a sample (for example, H calls in inbred samples, the presence of an unexpected diagnostic allele in “pure” sample, etc.). To address this issue, we reannotated the performance and diagnostic capability of all SNP markers. Reclustering for selected markers leads to better performance as measured by the level of consistent genotype calls between samples known to carry the same allele. Importantly, some identical (or related) samples genotyped using both the original and the new clustering parameters may have differing genotypes at some markers. Users should check whether these markers are annotated as reclustered in **Table S2.** In these cases, changes in genotypes are likely due to the analytical pipeline rather than biological reasons. In general, the genotypes generated with the updated pipeline should be preferred.

In the future, adding new samples may result in upgrading some markers from low performance to a higher performance tier. In addition, we will continue to manually curate the tier of markers with inconsistent annotations (for example, tier and diagnostic value) or subpar performance in specific samples. If the number of markers with annotation changes is significant, we will consider releasing a public update.

The report uses only Tier 1A markers to determine the primary and secondary backgrounds. The report uses all Tier 1A, 1B, 2A, and 2B markers annotated as diagnostic in the Backgrounds Detected (Diagnostic Alleles) section. The selection of markers from different Tiers in these different analyses reflects that Tier 1A markers are reliable across all genetic backgrounds for local and global genome analysis, and the additional four Tiers listed above do not falsely detect diagnostic alleles in any genetic background tested.

Our general advice to those planning to use MiniMUGA genotype data directly is to submit enough biological replicates from both sexes for the backgrounds of interest (we also recommend genotyping the corresponding F1s hybrids), and to use those genotypes to select markers that are both informative and have fully consistent genotypes in parentals and F1s. These will be mostly Tier 1A markers but some Tier 1B, 2A, and 2B may be selected and useful for specific needs.

Including the mitochondria and the Y Chromosome in the ideogram is a long overdue improvement that will easily alert users of congenic strains, among others, of potential errors or limitations in their mice. When an ideogram has regions where the primary and secondary backgrounds are highly fractured (having frequent changes of genetic background and/or zygosity over a short genomic distance), as shown in **Figure S4**, the selected genetic backgrounds are likely incorrect, or there are additional backgrounds present. Fracturing can also be localized, and the same conclusion applies. The report will provide a warning in the Summary section if fracturing is detected.

Diagnostic alleles are recent mutations that arose in an inbred line in the ancestors of one or more of its substrains. They were intentionally overrepresented in MiniMUGA to discriminate between closely related substrains. They are a unique feature of the MiniMUGA design in part because their inclusion necessitates prior knowledge of the whole genome sequence of the relevant substrain. Given their importance in genetic QC, the reliability of genotypes at diagnostic markers is worth considering. The vast majority (95%) of diagnostic SNPs are in the best performing Tier 1A. Manual curation has been employed to confirm that samples containing a diagnostic allele (in homozygosity and/or heterozygosity) are consistent with the presence of the substrain(s) indicated in the annotated diagnostic class. For diagnostic SNPs where the diagnostic allele is very rare or absent (in homozygosity and heterozygosity), the reliability of those genotypes is lower due to lack of precision in the clustering algorithm. As a rule, the reported presence of a substrain based on a single heterozygous call at a diagnostic SNP should be treated with caution.

An operational definition of diagnostic SNP is that the minor allele is only observed in the annotated substrains. The reliability of the diagnostic annotation depends on the size and genetic diversity of the outgroup of classical inbred strains in which the diagnostic allele is absent. In this analysis, the diagnostic outgroup used has at least 36 distinct classical inbred strains (27 strains in the common outgroup plus 9 strains with substrains) representing most genetic diversity in the classical inbred strains commonly used in mouse-based research (**Table S4**; Yang et al. 2011). Furthermore, each diagnostic allele is absent in an average of 87.6 (± 0.6 SD) classical inbred strains and substrains in our consensus set. (**Table S4**; Yang et al. 2011). Future addition of new substrain samples may lead to annotation change from fully diagnostic to partially diagnostic. The addition of new substrains may lead to new diagnostic class annotations.

In conclusion, the updates in the MiniMUGA pipeline reported here reduce the impact of under-performing SNP markers, increase the reliability of the reported genetic backgrounds, provide a better estimation of background contribution and level of inbreeding, expand the universe of samples for which a full report is generated, and provide new information including the potential presence of two additional construct and detailed diplotype intervals. We hope these changes are useful, and we welcome comments from the community regarding further enhancements. Non-expert users may want to take advantage of a recently released short video tutorial that provides a 10-minute guide on how to interpret a MiniMUGA sample report (https://www.med.unc.edu/mmrrc/).

## Data Availability and Software

All sample genotype and intensity data used in the supporting analyses and development of this manuscript have been archived at the University of North Carolina Dataverse (Blanchard 2024, https://doi.org/10.15139/S3/YQLDUJ)

## ACKNOWLEDGENTS

This work was supported in part by the following NIH grants Mutant Mouse Resource and Research Centers U42OD010924 (to TM), U24HG010100 (to LM, FPMV), R01ES029925 (to FPMV, RF, MS), P42ES031007 (to RF, FPMV), U19AI100625 (to FPMV, MF, MTH, RSB), R21DA052171 (NIDA to LT), U01AI149644 (to RSB), AI132130 (to CMS), AI157253 (MH and RSB), R01HL155986 (to TK), and P01AI132130 (to FPMV, MF, CS, LT). We thank the Systems Genetics Core Facility for access to the CC genotypes. The MiniMUGA genotyping array was developed and updated under service contracts to FPMV at UNC from Neogen Inc., Lincoln, NE. The authors have no conflict of interest to declare. None of the authors have a financial relationship with Neogen Inc. apart from the service contracts listed above. We are grateful for Neogen/Transnetyx for providing genotypes for consensus of several substrains from Janvier Labs.

## SUPPLEMENTARY MATERIAL

**Table S1**. List of samples used in this update of MiniMUGA. The table provides a randomly assigned ID, the laboratory source of the samples, the type of sample regarding the level of inbreeding, a reference for previously published samples, whether the sample was used in reclustering exercise, used in building the consensus genotypes for a given strain or serve as positive or negative control for the two new constructs added to the report.

**Table S2**: Marker annotation updates. This file includes marker annotations for the following fields:

- chromosome – Marker chromosomal location. Possible values are 1-19 autosomes, X and Y sex chromosomes, PAR pseudoautosomal region, or 0 for unmapped genetic construct probes.
- position – location in base pairs
- name – marker name
- tier_2022 – new SNP marker performance tier assignment. 1A, 2A, 1B, 2B, 1C, 2C, and 4 are possible values. Construct probes have no tier_2022 assignment.
- diagnostic – the list of substrains (or diagnostic class) for which this SNP is found to be diagnostic, or blank if it is not diagnostic.
- partial_diagnostic - the list of substrains for which this SNP is found to be partially diagnostic, or blank if it is not partially diagnostic.
- diagnostic_allele – the minor (diagnostic) allele at this diagnostic SNP.
- diagnostic_info – this field indicates which genetic construct this marker detects, or blank if it does not detect a construct (or detection has not been validated)
- positive_threshold – for markers that detect the same genetic construct, this value indicates the minimum threshold for reporting positive detection (presence), based on the sum intensity at these markers.
- negative_threshold – for markers that detect the same genetic construct, this value indicates the maximum threshold for reporting negative detection (absence), based on the sum intensity at these markers.
- duplication – This field indicates whether the marker probe appears to be duplicated in some samples in our performance sample set. This duplication may be the cause of a tier_2022 downgrade from A to B or C class. This field is still under development and not exhaustive.
- recluster – Indicates whether the Illumina genotype calling parameters (clusters) were modified in this update. Markers that are annotated as TRUE may generate different genotype calls for the same intensity values (or for the same sample) after this update.

**Table S3:** Genotype dump for consensus genotypes. This table includes consensus genotype calls for 237 inbred strains and substrains at 10,819 SNP markers.

**Table S4**. List of classical substrains and strains. The rows list all classical strains while the columns list the strain groups used for determination of diagnosticity of SNPs and an outgroup with all classical inbred strains without diagnostic SNPs. Only cells with numbers or N/A should be considered. Numbers in each cell are the number of diagnostic SNPs followed by the number of partially diagnostic SNPs for each substrain.

**Table S5.** 23 examples of improved estimation of the contribution of the primary and secondary backgrounds.

**Figure S1.** Tier 1B, 1C, 2A, 2B, 2C and 4.

**Figure S2.** Intensity of individual probes for the new constructs detected by MiniMUGA

**Figure S3.** Example ideograms of F1 hybrids of related substrains

**Figure S4.** Example of fracturing of ideograms.

